# Unlocking prophage potential: *In silico* and experimental analysis of a novel *Mycobacterium fortuitum* LysinB containing a peptidoglycan-binding domain

**DOI:** 10.1101/2024.02.15.580446

**Authors:** Ritam Das, Kanika Nadar, Ritu Arora, Urmi Bajpai

**Affiliations:** Faculty of Biological Sciences, Friedrich Schiller University, Jena-07737, Germany; Department of Life Science, Acharya Narendra Dev College, University of Delhi, Govindpuri, Kalkaji-110019, New Delhi, India; Department of Biomedical Science, Acharya Narendra Dev College, University of Delhi, Govindpuri, Kalkaji-110019, New Delhi, India

**Keywords:** Prophage, Endolysins, LysinB, Structure, Domains, Esterase activity

## Abstract

Endolysins are highly evolved bacteriophage-encoded lytic enzymes produced to damage the bacterial cell wall for phage progeny release. They offer promising potential as highly specific lytic proteins with a low chance of bacterial resistance. The diversity in lysin sequences and domain organization can be staggering. *In silico* analysis of bacteriophage and prophage genomes can help identify endolysins exhibiting unique features and high antibacterial activity, hence feeding the pipeline of narrow-spectrum protein antibiotics. Mycobacteriophage lysis cassettes mostly have two lytic enzymes, LysinA and LysinB. The enzyme LysinA targets peptidoglycan in the cell wall and possesses a modular architecture. LysinB typically contains a single domain and acts upon the mycolyl ester linkages in mycolyl-arabinogalactan-peptidoglycan (Payne *et al*., 2010). This study aimed to find novel LysinBs against *Mycobacterium fortuitum*. After a detailed *in silico* characterization of lysis cassettes from three *M. fortuitum* prophages, we chose to work on a LysinB (hereafter described as LysinB_MF) found in an incomplete prophage (phiE1336, 9.4 kb in strain E1336).

LysinB_MF showed low sequence similarity with any other endolysins in the database and formed a separate clade on phylogenetic analysis. LysinB_MF’s structure, extracted from the AlphaFold Protein Structure Database, demonstrated a modular architecture with two structurally distinct domains: a peptidoglycan-binding domain (PGBD) at the N-terminal and the characteristic alpha/beta hydrolase domain connected via a linker peptide. We found the alpha/beta hydrolase domain, which is the enzyme-active domain (EAD), contains the conserved Ser-Asp-His catalytic triad with a tunnel-like topology and forms intermolecular hydrogen bonds. The PGBD shows structural similarity to the cell-wall binding domain of an amidase from *Clostridium acetobutylicum,* hinting at its acquisition due to domain mobility. Our *in silico* electrostatic potential analysis suggested that PGBD might be essential to the enzyme activity. Based on our analysis, PGBD emerged as an integral constituent of enzymes with diverse functional properties and is predicted to be conserved cross-kingdom. Overall, this study highlights the importance of mining mycobacterial prophages as a novel endolysin source. It also provides unique insights into the diverse architecture of mycobacteriophage-encoded endolysins and the importance of functional domains for their catalytic activities.

## 1. Introduction

*Mycobacterium* infections are among the most debilitating, if not fatal (Gordon & Parish, 2018; Das *et al*., 2021). *Mycobacterium tuberculosis*, the causative agent of Tuberculosis, alone was the cause of 1.25 million deaths in the year 2023, replacing COVID-19 as the world’s leading cause of fatality from a single infectious agent (Goletti *et al*., 2025). Infections due to nontuberculous *Mycobacterium* (NTM) are also becoming an emerging issue of concern, and their prevalence is increasing worldwide. While *Mycobacterium fortuitum-*associated pulmonary diseases are the most prevalent (Khosravi *et al*., 2018), the bacterium also causes extra-pulmonary infections in humans, such as cutaneous and bone lesions, which are difficult to treat, chronic, occult, resistant, and recurrent (Sethi *et al*., 2014; Fraga *et al*., 2022).

Mycobacteriophages are viruses that specifically infect *Mycobacterium spp*. Most mycobacteriophages code for two endolysins: LysinA and LysinB, which target the bacterial cell wall and cause lysis of their host cells at the end of their life cycle (Payne *et al*., 2010; Payne & Hatfull, 2012). LysinA is a peptidoglycan hydrolase, and LysinB is a mycolylarabinogalactan esterase, which hydrolyzes the mycolyl ester linkages in mycolyl-arabinogalactan-peptidoglycan layer present in mycobacteria. Since the exogenous application of endolysins has been demonstrated to show an antibacterial effect, lysins have been explored as an alternative/combination therapy with antibiotics in several studies (Fraga *et al*., 2019; Abouhmad *et al*., 2020; Hurst-Hess *et al*., 2023; Singh *et al*., 2023). While the reports on structural and functional characterization of mycobacteriophage-encoded lysins (Payne *et al*., 2010; Eniyan *et al*., 2020; Gigante *et al*., 2021) have provided insights into their structural diversity, these lysins are relatively less reported when compared to phage lysins against Gram-negative and Gram-positive bacteria.

Besides the isolated mycobacteriophages, the genomes of nontuberculous *Mycobacterium* (NTM) that are enriched with prophages (Abad *et al*., 2023), could also serve as a promising source for diverse antibacterial proteins (Schmitz *et al*., 2011; Shah *et al*., 2023). However, these prophages are largely unannotated. In this study, we report a novel LysinB encoded by one such unique prophage in a strain of *M. fortuitum,* with three characteristic features: i) unlike the usual single-domain architecture of LysinB enzymes, LysinB_MF has a structurally distinct peptidoglycan binding domain (PGBD), ii) its enzyme active domain shows structural similarity to a hydrolase (amidase) of *Clostridium acetobutylicum,* and iii) PGBD might have a role in enzyme activity. As a first step towards understanding the molecular properties, we have characterized the lytic cassette of three *M. fortuitum* prophages, and carried out a detailed examination of LysinB_MF, showing the highest similarity with D29 LysinB, which we used as a bait sequence for searching LysinB sequences in *M. fortuitum* prophages. We have carried out an extensive sequence and structural analysis of lysis cassettes and LysinB_MF and found computational evidence of the influence of PGBD in its activity.

## 2. Results

### 2.1 *Mycobacterium* prophages as a source of novel lysin sequences

D29 and Ms6 LysB enzymes are the extensively studied mycolylarabinogalactan esterases with pre-clinical characterization (Payne & Hatfull, 2012; Gigante *et al*., 2017; Fraga *et al*., 2019; Hurst-Hess *et al*., 2023). However, to facilitate the growing interest in mycobacteriophage lysins to treat drug-resistant *Mycobacterium spp.,* more lysins need to be discovered and investigated to understand their biological features and to develop them as novel therapeutics. Non-tubercular mycobacteria (NTM) are reported to harbour a large proportion of prophages (Abad *et al*., 2023). We used D29 LysB (O64205) as a bait sequence and performed BLAST to find similar but not identical LysinB sequences against all the *M. fortuitum* genomes in the database. The workflow for the *in silico* identification of LysinB enzymes is represented in Figure 1A. The top three hits for LysinB were found to be encoded by prophages of *M. fortuitum* strains (Table 1). In this study, we have carried out the structural and functional characterization of a unique LysinB enzyme (mentioned as LysinB_MF) encoded by a prophage, called phiE1336, present in the sequenced strain E1336 of *Mycobacterium fortuitum* (tax id: 1766). A detailed characterization of this prophage has been done in the later sections.

**Figure 1:**
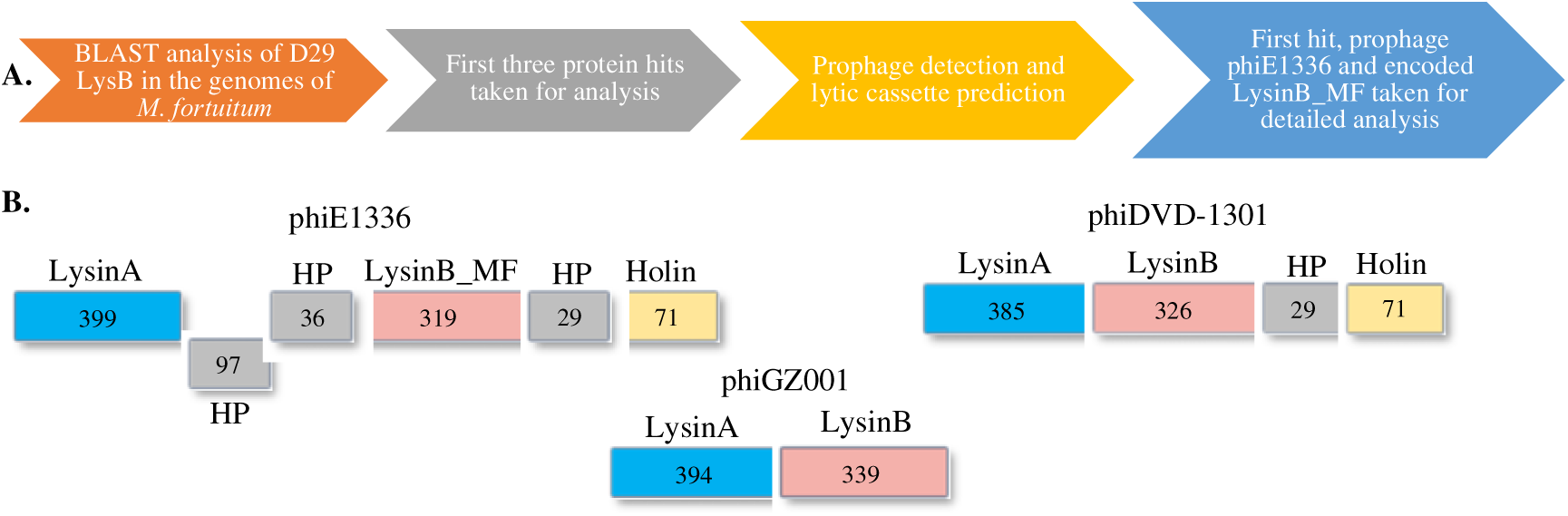
Analysis of prophages in *M. fortuitum* strains. **A)** The workflow of prophage detection and characterization of the lytic regions. **B)** Lytic cassette of prophages phiE1336, phiDVD-1301 and phiGZ001.

**Table 1:** Information on M. fortuitum prophages identified and characterized in the study.

| S. no. | Name | Size (kb) | Completeness | GC% | Cluster similarity | *M. fortuitum* lysogen |
| --- | --- | --- | --- | --- | --- | --- |
| 1 | phiE1336 | 9.4 | Incomplete | 62 | - | NZ_LZSJ01000108.1 |
| 2 | phiDVD-1301 | 39.6 | Intact | 62.8 | P | NZ_JARUNL010000131.1 |
| 3 | phiGZ001 | 38.9 | Intact | 63.4 | F1 | NZ_CP107719.1 |

We found the prophage phiE1336 genome to be incomplete (9.4 kb), however, its lysis cassette was intact. It constituted LysinA, three small hypothetical proteins, LysinB and Holin (Figure 1B). LysinB_MF, with an overall identity of 49% with D29 LysinB, is a peptidoglycan-binding domain-containing protein (WP_064866887.1). The second hit, with an overall identity of 48%, was a hypothetical protein (HP) (WP_278289780.1) from *M. fortuitum* strain DVD-1301 (NZ_JARUNL010000131.1). Reannotation of the genome and PHASTER prediction indicated the protein to be a LysinB, encoded by an intact prophage (hereby called phiDVD-1301). This prophage genome size was 39.6 kb with a GC content of 62.86%, similar to what is observed for other mycobacteriophages. Nucleotide BLAST analysis of the genome indicated phiDVD-1301 to have approximately 80% sequence identity with cluster P phages Tortellini (KX648391), Phayonce (KR080195) and Purky (MN096355). Annotation demonstrated the presence of a lytic cassette (Figure 1B) with LysinA (385 amino acids) as the largest protein-coding gene. LysinB (326 amino acids) from phiDVD-1301 showed 48% identity with D29 LysB, however, domain predictions for this enzyme remained unsuccessful. The third hit (WP_269975983.1) showing 36% sequence identity with D29 LysB was a hypothetical protein (HP), encoded by an *M. fortuitum* strain MF GZ001 (NZ_CP107719.1) prophage. This prophage (hereby called phiGZ001) has a size of 38.9 kb with distinct attL (CACACTGTTCTTC) and attR (CACACTGTTCTTC) sites recognized by serine integrases (Suzuki *et al*., 2020). Nucleotide BLAST analysis of the genome indicated phiGZ001 to have approximately 86% sequence identity with cluster F1 phages XFactor (KT281795), Tootsieroll (MZ747519) and Blexus (MN586012). Sequence reannotation and domain analysis of the identified HP (WP_269975983.1) predicted an alpha/beta hydrolase domain. In phiGZ001, we did not find a holin gene in the lysis cassette (Figure 1B). We speculate this prophage either lyses the host independent of holin (Eniyan *et al*., 2020) or the lytic cassette is incomplete.

### 2.2 The incomplete prophage in the *M. fortuitum* strain E1336

PHAST analysis predicted the presence of an incomplete genome of prophage phiE1336. Its size is only 9.4 kb (Figure 2A) but the GC content (62%) is comparable to the host (Ho *et al*., 2012). Mycobacteriophages are grouped into clusters based on the similarities and differences in their nucleotide sequences (Hatfull, 2022). Currently, 32 clusters of mycobacteriophages have been defined, and the unclassified phages are denoted as singletons (https://phagesdb.org/). We performed nucleotide BLAST analysis to identify the possible cluster of phiE1336 and to predict phages identical to it. None of the whole genome sequences in the database showed more than 10% query coverage, a trend similar to singletons. However, considering the short genomic fragment of phiE1336, it is difficult to state conclusively whether it belongs among the singleton phages or to a cluster.

**Figure 2:**
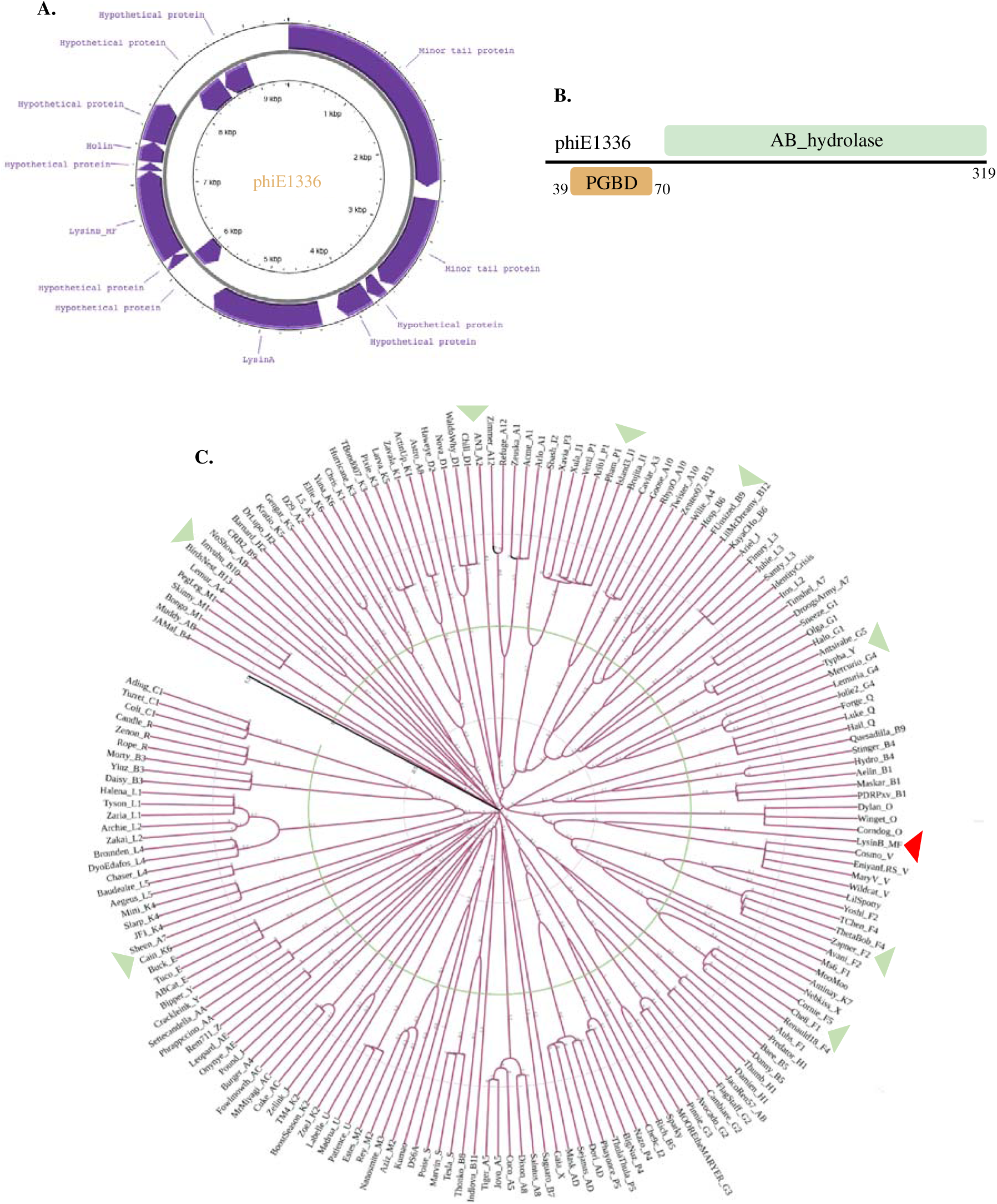
Characterization of phiE1366 and the encoded LysinB_MF. **A)** phiE1336 has a genome size of 9.4 kb with a GC content of 62%. The genome consists of 13 ORFs, which encode for a few structural proteins and proteins of the lytic cassette **B)** Domain architecture of LysinB_MF from phiE1336 demonstrates distinct PGBD and EAD. **C)** Phylogenetic analysis of LysinB_MF to understand its evolutionary relationship. Representative sequences of LysinBs from all sub-clusters and singletons were taken for analysis. Phylogeny analysis displayed LysinB_MF forming its separate clade (in red arrow), which branched from the neighbouring Cluster V LysinB at some point. Green arrows show the clades disparate with the evolutionary relationships of mycobacteriophage clusters.

### 2.3 The lytic cassette in prophage phiE1336

We extracted the 9.4 kb fragment and performed annotation using RAST, which predicted it to have 13 ORFs (Figure 2A). Among the predicted ORFs, six could be assigned putative functions, which included lytic and structural proteins (Table S1). The lytic cassette begins with LysinA (44.3 kDa), a peptidoglycan hydrolase consisting of an amidase domain spanning from 171–317 and an overlapping peptidoglycan recognition protein (PGRP) region from 183–317 amino acids (Figure 2B). Co-occurrence of domains is common in phage lysins (Payne & Hatfull, 2012; Hatfull, 2018). The amidase domain targets the N-acetylmuramoyl-L-alanine linkages, which connect the sugar backbone to the peptide crosslink (Payne & Hatfull, 2012). BLAST analysis showed phiE1336 LysinA to share 71.9% identity with LysinA (gp24) from F1 sub-cluster mycobacteriophage Koella (MH316564) and 70.5% identity with LysinA (gp69) from O cluster phage Smooch (MN428052). Domain construction of lysins from these phages is similar and constitutes the active site of the amidase with an overlaying peptidoglycan recognition protein (PGRP) region.

The two small HPs between LysinA and LysinB of phiE1336 (Figure 1B) are of size 10.6 kDa and 3.9 kDa and do not possess a recognizable domain. Neither of the HPs shows any significant similarity to other sequences in the database. This arrangement of the lytic cassette, with HPs between Lysin A and LysinB, appears unique for phiE1336 among the other prophages characterized in this study (Figure 1B). The other reported mycobacterium prophages, such as phiMAV1 and MabR, constitute lytic regions with LysinA and LysinB present together (Payne *et al*., 2010; Cote *et al*., 2022).

Upstream to the two HPs in the cassette are the genes encoding for mycolylarabinogalactan esterase (LysinB_MF), another small HP (3.1 kDa), and holin (Figure 2B). LysinB_MF is a multidomain enzyme with a molecular weight of 35.3 kDa. While we could not precisely predict the function of the 3.1 kDa protein, BLAST analysis showed similar proteins to have an ATP binding activity. The holin gene encodes for a protein of size 7.4 kDa, which shares 61.1% identity with the holin from cluster I2 mycobacteriophage Che9c. Comparable to other phage-encoded holin, this, too, consists of two transmembrane helices, which could destabilize the membrane proton motive force of the host in a time-dependent manner (Pollenz *et al*., 2022).

### 2.4 LysinB_MF

Although the information on mycobacteriophage-encoded mycolylarabinogalactan esterase (LysinB) has been discussed earlier in several studies (Payne *et al*., 2010; Korany *et al*., 2019; Abouhmad *et al*., 2020), reports on the characterization of a multidomain LysinB is limited (Gigante *et al*., 2021). We found LysinB_MF to be a modular enzyme composed of two structurally distinct domains connected by a linker peptide (Figure 2B). The enzyme (319 amino acids) is comprised of two functionally defined regions: PGBD and the alpha/beta hydrolase domain (Figure S1). The PGBD is present at the N-terminal region of LysinB_MF and spans from amino acids 39–70, and a linker of 14 amino acids connects it to the catalytic enzyme-active domain (EAD) (Figure 2B). The EAD, like most mycolylarabinogalactan esterases, is comprised of an alpha/beta hydrolase domain that spans right after the linker until the C-terminal end, covering a substantial part of the enzyme. Through BLAST analysis, we found LysinB from a V cluster mycobacteriophage Cosmo (KP027195) to be the closest predicted homologue of LysinB_MF. The sequences showed a high query coverage of 98% and 56.43% identity (e-value 3e-116). Besides this, we could not find duplicate or identical proteins to LysinB_MF in the database. Therefore, to examine the evolutionary relationship of LysinB_MF, we performed the phylogenetic analysis using LysinB sequences from singletons and mycobacteriophages belonging to all the sub-clusters (Figure 2C). There were 187 sequences in the repository file prepared with at least one sequence and a maximum of three. The multi-FASTA file was subjected to sequence alignment and then processed using the iTOL to generate a rooted phylogenetic tree based on the maximum likelihood method.

Interestingly, we found a separate clade for LysinB_MF, as observed for LysinBs from singleton phages (Figure 2C). The LysinB_MF sequence appeared to have branched from the cluster V endolysins and had a branch length of 0.79. A PGBD in these LysinBs remains unannotated or lacking, although conserved residues in the N-terminal region were observed in the multiple sequence file. It appears Cluster V LysinBs and LysinB_MF may have had a common ancestor at some point; however, they diverged into forming their clade, and LysinB_MF retained the PGBD function.

Though the phylogenetic analysis gave us some clue about the evolutionary relationship of LysinB_MF, what was interesting to note was the unconventional grouping of endolysins from different sub-clusters in the same clade (shown in green arrows, Figure 2C). Phylogenetic analysis of tape-measure proteins in mycobacteriophage genomes distinctly shows separate clades, called clusters or even specific sub-clusters (Jacobs-Sera *et al*., 2012). Phages belonging to the same cluster are grouped into the same clade. However, while studying LysinB, we observed sequences belonging to entirely different clusters, if not sub-clusters, forming a common sub-clade and demonstrating the possibility of a common ancestor at some point in the evolutionary history. LysinB from clusters A7 and K6; F1, F4 and F5; B6 and B12; P1, P3 and I1, respectively, were present in the same clade (Figure 2C). In some instances, the evolution of LysinB appeared to be independent of the evolution of mycobacteriophages. Mycobacteriophage-encoded LysinBs, therefore, emerge as enzymes with a complex evolutionary pattern, and their lineage requires a greater analysis.

### 2.5 LysinB_MF has a complex enzyme active domain (EAD)

To perform the structural and functional analysis of LysinB_MF, we attempted to characterize the 3D model of the enzyme available in the AlphaFold Protein Structure Database (AFDB: A0A1A0NU50) and carried out *in vitro* assays of the purified protein. The structure of this enzyme was available as a peptidoglycan-binding protein derived from the *M. fortuitum* strain described earlier. The protein structure showed a high model confidence (pLDDT>90) and was taken forward for further analysis (Figure 3A and 3B).

**Figure 3:**
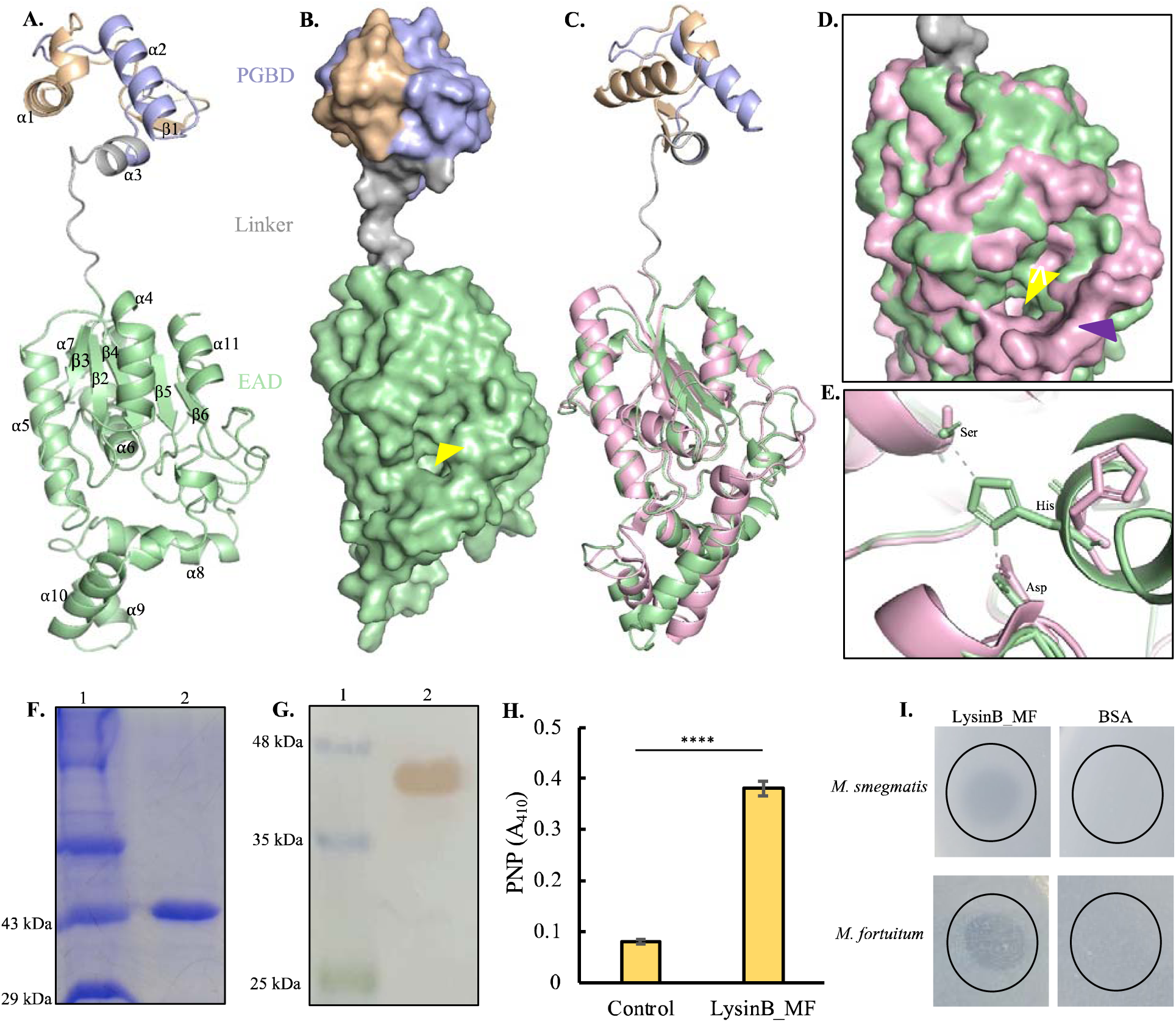
Structural and functional analysis of LysinB_MF. **A)** Three-dimensional (3D) model of LysinB_MF demonstrates a multi-domain structure comprising two structurally distinct domains: the PGBD (39-70), a short linker region followed by the EAD, an lz/β hydrolase (85-317). **B)** The surface model of LysinB_MF demonstrates a bulky C-terminal end with a distinct active-site groove formed with residues Ser-Asp-His. The groove of the EAD, indicated with a yellow arrow, displays a tunnel-like topology. **C)** LysinB_MF can be superimposed on D29 LysB (in pink) with an overall RMSD of 0.56. Structural alignment of the two endolysins shows similar active-site arrangement, although PGBD is absent in D29 Lysin B. **D)** Active-site conformation in LysinB_MF and D29 LysinB. LysinB_MF shows a narrow but deeper topology (shown with a yellow arrow). D29 LysB displays a larger edge, having a funnel-like conformation (shown with a purple arrow). **E)** Active site residues in LysinB_MF (Ser156-Asp236-His305) and D29 LysB (S82-Asp166-His240). **F)** SDS-PAGE (12%) analysis of purified LysinB_MF. Lane 1: protein molecular mass standards. Lane 2: purified N-terminal His-tagged LysinB_MF, molecular weight 43 kDa (accurate) and 37.5 kDa (average). **G)** Immunoblot of N-terminal His-tagged LysinB_MF. Lane 1: protein molecular mass standards. Lane 2: N-terminal His-tagged LysinB_MF. **H)** Esterase activity of LysinB_MF determined by the PNP release assay (n=9) at 1 µg of enzyme preparation. Asterisks indicate significant differences estimated by Student’s t-test (****: p < 0.0001). **I)** Spot test for intrinsic activity of LysinB_MF (10 µg) against *M. smegmatis* and *M. fortuitum*. BSA (10 µg) was used as a negative control.

The EAD of LysinB_MF comprises the [Z/β sandwich belonging to the [Z/β hydrolase family and contains the catalytic triad Ser-Asp-His. This domain spanned from amino acid 85–317, covering a significant portion of the enzyme and demonstrated a highly twisted structure consisting of a C-terminal linker region from 246-310. The active site of LysinB_MF displayed four β-strands (β2–β5), surrounded from all sides by [Z-helices ([Z4-[Z6). A surface model of the enzyme demonstrated the active-site groove with a tunnel-like topology (Figure 3B). A similar topology is observed in D29 LysB, which serves as a binding pocket for substrates such as several *p-*NP derivatives, trehalose monomycolate and mAGP from mycobacterial cell wall (Payne *et al*., 2010). A DALI search with high confidence (DALI Z-score 29.7) identified the structural relationship of LysinB_MF with the tertiary structure of D29 LysB (PDB accession 3HC7). The two esterases could be superimposed with an overall RMSD of 0.56 (153 C^l1^ atoms) (Figure 3C). Both the structures displayed similar β-sheet conformation around the active site surrounded by the helices. Given the high structural similarity observed between LysinB_MF and D29 LysinB, the topology of D29 LysB’s binding pocket, unlike LysinB_MF, showed an edge with a wider circumference and a funnel-like appearance consisting of key active site residues (Figure 3D and 3E). Furthermore, a bird’s eye view and analysis of the conserved active-site residues (Ser82, Asp166 & His240 in D29 LysB (Payne *et al*., 2010) and Ser156, Asp236 & His305 in LysinB_MF) of these enzymes displayed notable differences (Figure 3E). Unlike their counterparts, the residues Asp166 in D29 LysB and His305 in LysinB_MF were part of alpha-helices in these enzymes, which added to the differences in their groove topology. Also, unlike D29 LysB, the active-site residue His305 in LysinB_MF was found to form polar contacts with residues Ser156 and Asp236 (Figure 3E). These bonds are known to provide stability by acting as electrostatic attractive forces which influence the charge redistribution in the enzyme and are responsible for the uptake of electrons (pull) from the active site towards the amino acid residues (Qiu *et al*., 2018). Also, in the case of serine proteases with similar Ser-Asp-His catalytic triads, the removal of hydrogen bonds has been shown to result in the partial release of the active-site residues in molecular dynamic simulations and a decrease in the overall stability of the transition states, which affects the enzyme’s turnover number (*k*_cat_) (Zheng *et al*., 2006). In the *in vitro* assay with the purified LysinB_MF (Figures 3F and 3G), we observed significantly high esterase activity of the enzyme as evident by PNP release compared to the control (buffer without enzyme) (Figure 3H). Using a spot test, we also found LysinB_MF to cause lysis of non-pathogenic *M. smegmatis* and pathogenic *M. fortuitum* cells (Figure 3I). We speculate that the predicted differences in groove topology and improved stability of the active site core might have a role in the enzyme activity.

The presence of such a potent lytic enzyme encoded by a truncated prophage, incapable of completing its life cycle to form and release phage progeny, is intriguing We hypothesise the probable reason behind *M. fortuitum* strain retaining this supposedly toxic protein could be either a chance event or the bacterial host uses it as an autolysin, keeping the expression of this enzyme tightly regulated. Future experiments employing genome engineering to create deletion mutants and analysis of the engineered bacterial strain could help validate these hypotheses.

### 2.6 The PGBD of LysinB_MF is structurally similar to an amidase

In most of the mycolylarabinogalactan esterases reported earlier (Fraga *et al*., 2019; Abouhmad *et al*., 2020; Eniyan *et al*., 2020; Singh *et al*., 2023), an extended N-terminal region consisting of a PGBD hasn’t been reported. It is a rare feature that has been observed in Ms6 LysinB (Gigante *et al*., 2021). Region 7–71 in LysinB_MF was interpreted as a PGBD-like superfamily, with region 39–70 predicted to be the functional PGBD (highlighted in light blue in Figure 2A). Only a single β-sheet in this region is present in the enzyme’s N-terminal end, and the rest of the structure appeared to be highly twisted, dominated by three [Z-helices. The annotated PGBD is comprised of two [Z helices ([Z2 and [Z3), and it tapers to form the linker region connecting to the EAD. The multiple sequence alignment and structural analysis of LysinB_MF demonstrated several conserved residues in this region (Figure 4), which are also found in LysinBs without the N-terminal PGBD. The amino acids with the highest score in the conservation index are mostly buried in the enzyme and could be involved in the functional properties of the PGBD (Figures 4A-4C).

**Figure 4:**
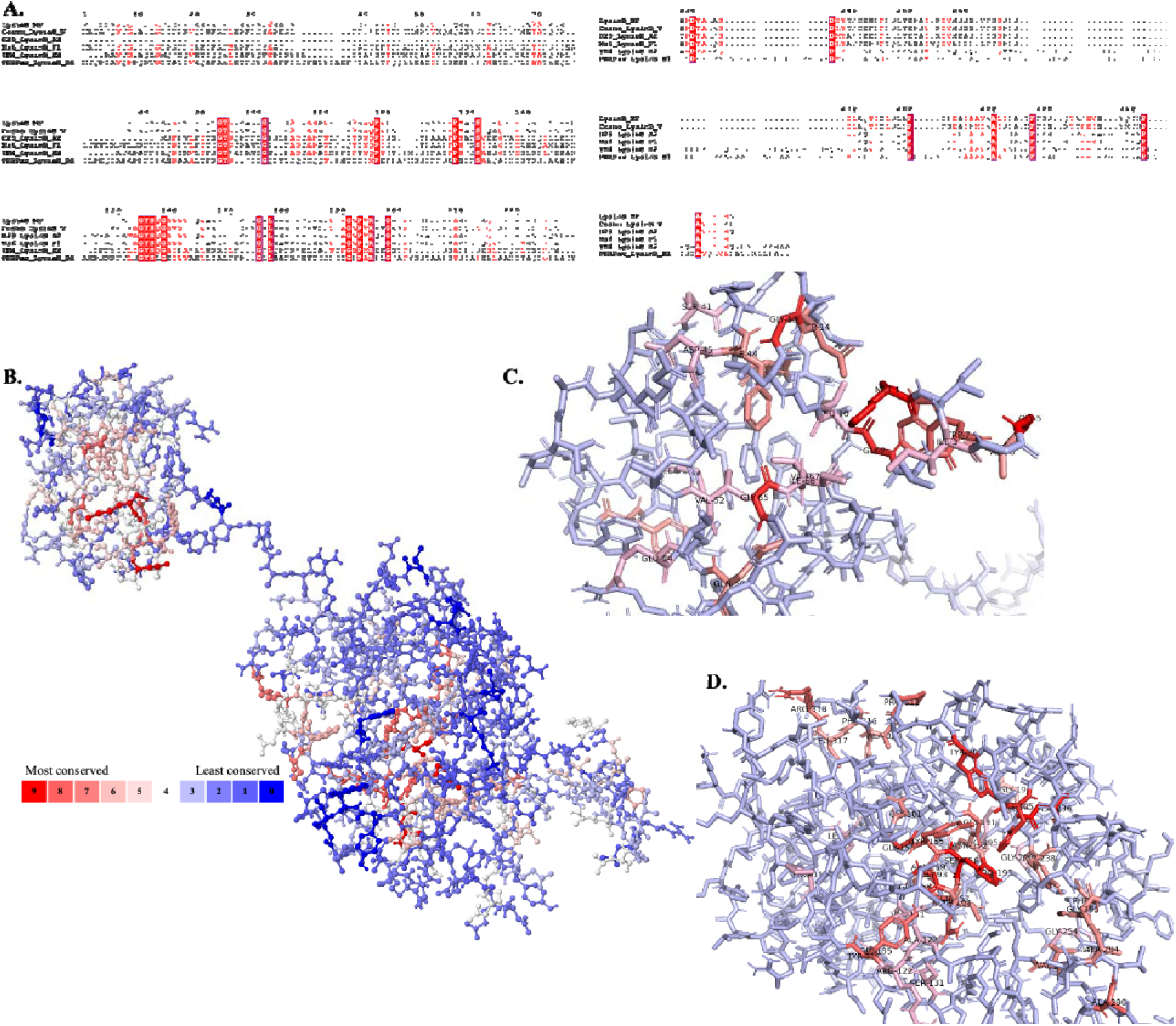
Conserved residues in LysinB_MF. **A)** Multiple sequence alignment of LysinB enzymes from various clusters demonstrate key residues in PGBD and EAD. The EAD displays the conserved active-site residues Ser-Asp-His and the GXP motif. **B)** Analysis using POLYVIEW-MM identified and mapped conserved residues in the tertiary structure of LysinB_MF. A scale of 0-9 is used to quantify the conservation index, where 0 indicates the least conserved and nine is the most. Exposed residues in the structure of LysinB_MF are found to be least conserved among all others, while most of the highly conserved residues are found buried. **C)** Bird’s eye view of PGBD displays conserved amino acids. Ser41, Tyr43 Phe44, Asp45, Val52, Glu54, Gln56, Gly66 and Ile67 are a few highly conserved residues in this domain found both in sequence alignment and in structural analysis. **D)** Bird’s eye view of EAD displays conserved amino acids. The active site Ser156, Asp236 & His305 and Gly191 & Pro193 from the GXP motif were highly conserved in LysinBs.

Considering the unavailability of structural information about the PGBD in a mycobacteriophage-encoded LysinB and failure to acquire the same through structural similarity analysis based on the native structure of LysinB_MF, we separately built the PGBD and analyzed it further. The tertiary structure of PGBD was prepared using LysinB_MF as a template, and it had a QMEANDisCo Global score of 0.63 ± 0.11, which ensured high quality (Figure S2A). We then performed structural similarity prediction using DALI, which predicted 20 possible neighbours, and these were visually sorted based on their Z-score. Interestingly, the top scoring structure (Z-score from DALI 9.5) found similar to the PGBD of LysinB_MF was an enzyme with amidase activity (PDB accession code 4XXT). This zinc-dependent amidase is a PGBD-containing protein from *Clostridium acetobutylicum* (ATCC 824). To analyze the structural similarity to PGBD of LysinB_MF, we downloaded the protein 3D file from the PDB database and carried out structural alignment using PyMOL. The PGBDs from the amidase and LysinB_MF could be superpositioned with an overall RMSD as low as 0.46, even though the sequences do not show identical residues (Figure S2B). Also, amidases (or lysozyme-like domains) possessing a peptidoglycan-binding and hydrolyzing function are usually the features of mycobacteriophage LysinA enzymes. Due to the unavailability of the 3D structure of a LysinA in the PDB database, we could not examine the structural similarity of PGBD specifically to a mycobacteriophage-encoded N-acetylmuramoyl-L-alanine amidase. However, it is noteworthy that structural similarity of PGBD of LysinB_MF with high confidence was achieved with an amidase from a bacterial species. We speculate that domain mobility could be a reason behind the generation of modular proteins, and they play a crucial role in their functioning (Basu *et al*., 2009).

### 2.7 PGBD is perhaps essential for the esterase activity of LysinB_MF

To understand the role of PGBD in the activity of LysinB_MF, we computationally generated its truncated version (LysinB_MFΔPGBD), lacking the N-terminal domain (Figures 5A and 5B). The truncated structure was built based on the native LysinB_MF as a template using SWISS-MODEL (0.70 ± 0.05). Using eF-surf, a comparative analysis of the native and the truncated structures indicated a change in the electrostatic potential of the active site. The catalytic tunnel appeared more electronegative in LysinB_MFΔPGBD than its native structure (Figures 5C and 5D). A change in electrostatic potential could impact the interactions between an enzyme and its substrate and, hence, the enzyme’s activity. The importance of electrostatics in enzyme-mediated catalysis has been demonstrated for amylases where a change in the charge near the active site residues resulted in an enzyme with drastically reduced activity (Nielsen *et al*., 1999).

**Figure 5:**
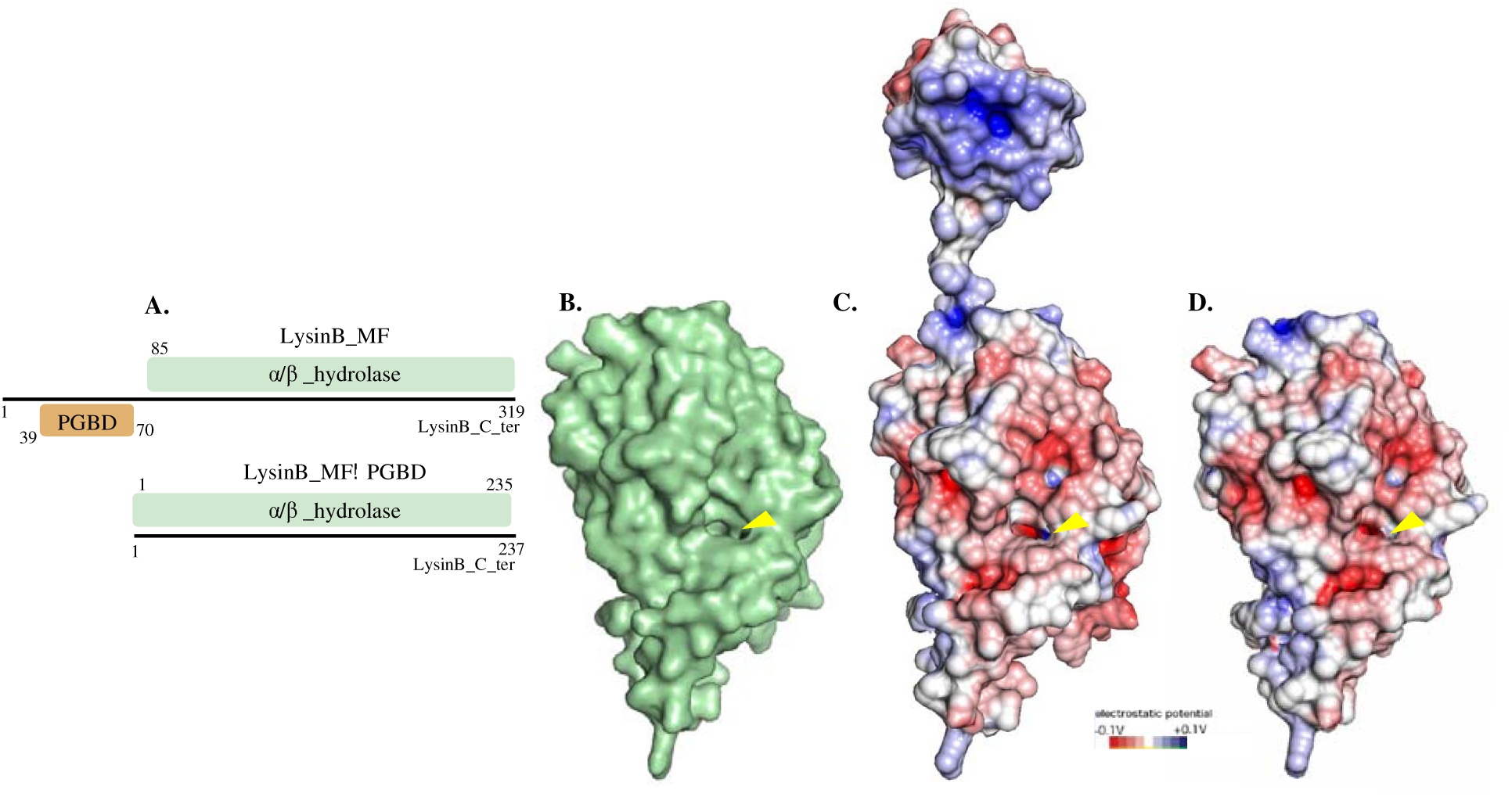
PGBD and active-site electrostatics of LysinB_MF. **A)** Domain architecture and comparison of native LysinB_MF and truncated LysinB_MFLiPGBD (EAD). **B)** The structure of LysinB_MF without the PGBD (LysinB_MFLiPGBD) was built using the native structure as a template. LysinB_MFLiPGBD structure comprises the linker region (in grey) and the EAD (in green). **C-D)** Comparison of the intact LysinB_MF and LysinB_MFLiPGBD demonstrates a change in electrostatic potential in the active site (indicated with a yellow arrow).

### 2.8 Conservation of PGBD across different kingdoms

Through *in silico* protein truncation, we predicted the integral role of PGBD in LysinB_MF activity (Figures 5H-5I). However, this feature is not constant in all other endolysins. While in the case of bacteriophage PSA, removing the cell-wall binding domain of endolysin demonstrated a similar decrease in enzymatic activity (Korndorfer *et al*., 2006), the lytic activity of a *Bacillus anthracis* prophage endolysin was enhanced when the C-terminal cell-wall binding domain was removed (Low *et al*., 2005). PGBD appears to have diverse functionality; however, a proper understanding of the conservation and importance of PGBD in all forms of life is limited. To comprehend the importance of this functional domain in other organisms, we analyzed all the sequences present in the UniProt database against the Pfam entry for the PGBD (PF01471). There are approximately 122,000 protein sequences consisting of this domain. From our curated analysis of these sequences based on TaxId from different organisms, we found this domain is conserved across all kingdoms (Figure 6). Among the prokaryotes that we analysed in this study, PF01471 was predicted to be the highest in *E. coli* (Figure 6), where it is present in LD-transpeptidases, which are involved in cell wall adaptation to stress and for providing outer membrane stability (Aliashkevich & Cava, 2022). In *M. fortuitum*, PF01471 was found chiefly in enzymes with a PG hydrolase activity. In the evolutionary Tree of Life in higher kingdoms, with *Arabidopsis thaliana* as an example, our curated search predicted 27 UniProt protein matches against PF01471 (Figure 6). Most of the matched proteins in this species were found to be zinc dependent endo peptidases or matrix metalloproteinases, which are involved in root development and water uptake (Mishra *et al*., 2021). In higher animals, including humans, we received 54 UniProt protein matches against PF01471, which, notably, was higher than most prokaryotes analyzed in this study (Figure 6). In humans, a diverse group of essential enzymes possess a PGBD, including matrix metalloproteinases (also observed in *A. thaliana*), enzymes involved in the degradation of extracellular matrix and development of human bones such as Stromelysin-2 (Bord *et al*., 1998) and collagenases, which are implicated in the turnover of connective tissue matrix constituents (Knauper *et al*., 1996). The cross-kingdom conservation of the short PGBD observed here would require a deeper analysis and is beyond the scope of our study.

**Figure 6:**
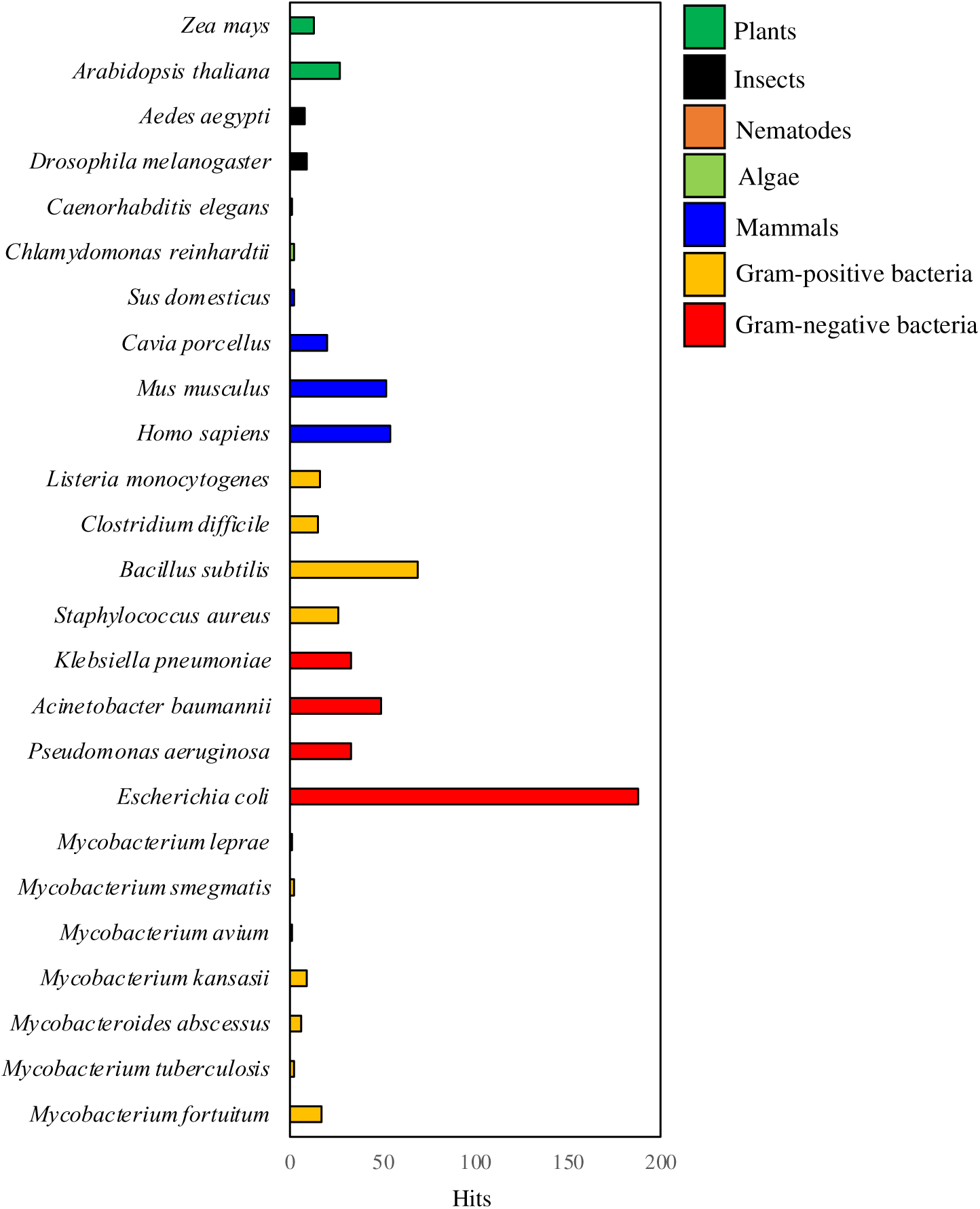
Cross-kingdom conservation of a PGBD. The UniProt database was screened against the Pfam ID of PGBD (PF01471), which resulted in approximately 122,000 protein hits with this domain. The curated database of hits was then individually screened against TaxIDs from various organisms across all kingdoms.

## 3. Conclusion

We conducted an *in silico* analysis of three prophages in *M. fortuitum*, a medically important NTM. The analysis includes extraction, annotation and sequence characterization of prophages, followed by the structural analysis of the lysis cassettes. In this study, a detailed analysis of one of the three prophages, ‘phiE1336’ and its encoded LysinB (LysinB_MF) was done. Though LysinB_MF is encoded by an incomplete prophage, which should be incapable of forming phage progeny on induction, it’s *in vitro* esterase activity is high. Investigating whether this active, prophage-derived LysinB enzyme has a role in the host *M. fortuitum* life cycle will be worthwhile. Besides the catalytic α/β hydrolase domain and a C-terminal linker region, LysinB_MF has an extended N-terminal, containing a peptidoglycan-binding domain. A conservation analysis predicted this domain in diverse enzymes encoded by several organisms. These findings underscore the critical importance of PGBDs in bacterial and viral enzymatic processes, their potential for functional specialization, and their evolutionary significance.

## 4. Materials and Methods

### 4.1 *In silico* identification of LysinB_MF and characterization

Lysin B from D29 (GenBank: AAC18452.1) was blasted (Altschul *et al*., 1990) against all the genomes in the NCBI database for *M. fortuitum* (tax id: 1766), possibly harbouring prophages. The *M. fortuitum* genome with the top lysin hits (LysinB_MF) was subjected to PHAST (Zhou *et al*., 2011) and to annotate and fix frameshifts in these genomes, the RAST server was used. The CGview tool (Grant & Stothard, 2008) was used to build the genome map of the first hit, phiE1336. Cluster-wise sequences of LysinB were downloaded from the Actinobacteriophage Database (https://phagesdb.org/), and domains of those lysins were predicted using InterProScan 5 (Quevillon *et al*., 2005). ClustalW (Thompson *et al*., 1994) and ESPript (https://espript.ibcp.fr) (Robert & Gouet, 2014) were used to generate alignment files which were then used to create phylogenetic trees using Interactive Tree Of Life (iTOL) v5 (Letunic & Bork, 2021). The tertiary structure of LysinB_MF was downloaded from the AlphaFold Protein Structure Database (Varadi *et al*., 2023) and visualized using PyMOL (https://pymol.org/2/). SWISS-MODEL was used to generate structures for truncated enzyme based on the AlphaFold predicted structure of LysinB_MF (Waterhouse *et al*., 2018).The POLYVIEW-MM tool was used to predict conserved residues in the tertiary structure of LysinB_MF (Porollo & Meller, 2010). The DALI server was used to find structures similar to the LysinB_MF (Holm, 2022) and eF-surf was used to predict the electrostatic potential of the enzyme (Nielsen *et al*., 1999). Conservation analysis was done by analyzing the Pfam entry of PGBD (PF01471) against all the proteins available in the UniProt database (https://www.uniprot.org/). TaxIDs were used to detect the presence of this domain across several species using the InterPro classification of protein family website (https://www.ebi.ac.uk/interpro/).

### 4.2 Cloning, overexpression and purification of recombinant LysinB_MF

The LysinB_MF gene was synthesized and cloned in the pET28a vector by Biomatik Corporation, Kitchener, Ontario. The recombinant construct pET28a-LysinB_MF was overexpressed in BL21(DE3) strain of *E. coli*. Induction conditions for soluble expression included 0.1 mM IPTG and overnight incubation at 16 °C. The induced culture pellet was collected by centrifugation at 8,000 rpm for ten minutes at 4°C and sonicated at 90% amplitude with 30 seconds each on or off pulse for ten cycles (Eniyan *et al*., 2020; Das & Bajpai, 2023). The cell lysate was centrifuged at 13,000 rpm for 45 minutes at 4°C, and the supernatant was considered the soluble fraction.

Recombinant LysinB_MF was purified by Ni-NTA affinity chromatography. Briefly, the soluble fraction containing the His_6_-tagged protein was kept for binding with the Ni-NTA matrix for 2 hours at 4 °C, and the bound resin was packed in a column. After packing and discarding the flow-through, the column was gradient-washed with buffer containing 20-40 mM imidazole and the bound protein was eluted at 200-250 mM imidazole concentrations. The eluted fractions were analyzed using 12% SDS-PAGE (Laemmli, 1970) and the purified protein was dialyzed with a buffer containing 50 mM Tris (pH 7.9) and 150 mM NaCl. The concentration of purified protein was estimated by Bradford assay (Bradford, 1976). For immunoblotting, the protein in the gel was transferred to a PVDF membrane using a Mini-PROTEAN Tetra Cell (Bio-Rad, United States). After blocking the membrane with 3% BSA at 4°C overnight, the membrane was incubated with mouse anti-His6 antiserum (Santacruz Biotech, USA) for 4 hours at a dilution of 1:2000. The membrane was washed once with PBS and twice with PBS-Tween, followed by incubation with IgG HRP-conjugated antibody at a dilution of 1:10000 for 45 minutes at 4 °C. The blot was developed by using 3, 3’-Diaminobenzidine (DAB-H_2_O_2_) as the substrate.

### 4.3 Assay for the esterase activity of LysinB

The micro-titre plate-based p-nitrophenol butyrate (pNPB) esterase assay (Fraga *et al*., 2019) was performed to estimate the activity of LysinB_MF. The enzyme reaction included 1 µg of enzyme 10 mM pNPB, and 25 mM Tris buffer (pH 7.2) in a final reaction volume of 200 µl. Incubation was done at 37°C for 30 minutes. Absorbance at 410 nm was measured to determine the release of p-nitrophenol (PNP). In the control reaction, enzyme buffer was added in place of the enzyme. The experiments were done thrice in three sets, and the absorbance value (A_410_) was plotted as mean ± SD.

### 4.4 Spot Test

To examine the antimycobacterial activity of LysinB_MF, the spot test on bacterial lawns was performed. Briefly, *Mycobacterium smegmatis* strain mc^2^155 and *M. fortuitum* (NIHJ 1615) were grown for 48 hours at 37°C using Middlebrook 7H9 broth (HiMedia, India), supplemented with 0.4% glycerol. Bacterial lawn in 7H10 Middlebrook agar plates containing 0.05% Tween 80 plates were prepared by spreading 0.5 mL of *M. smegmati*s/*M. fortuitum* using 7H10 soft agar. LysinB_MF (10 µg) was added to the prepared lawn as spots and the plates were incubated at 37°C and observed for the clear zones at the site of application.

## Acknowledgement

We thank the Science and Engineering Research Board (SERB), Department of Science and Technology (DST), India, for supporting the project (SERB PROJECT EMR/2017/004051). Acharya Narendra Dev College (ANDC), University of Delhi, is acknowledged for providing the infrastructural facilities and ELITE fellowship to RD. KN received the Innovation in Science Pursuit for Inspired Research (INSPIRE) fellowship from the Department of Science and Technology (DST), India; RA received the Senior Research Fellowship (SRF) from the University Grants Commission (UGC), New Delhi, India. The graphical abstract was created using BioRender.com (https://biorender.com).

## Author Contribution

RD: Data curation, Software, Visualization, Investigation, Validation, Writing-original draft. KN: Data curation, Experimental Investigation, Visualization, Validation. RA: Primer-designing, Data curation, Visualization. UB: Conceptualization, Methodology, Experimental Investigation, Supervision, Validation, Writing– review & editing.

## Availability of data and materials

This published article and its supplementary information files include all data generated or analyzed during this study.

## Ethics approval and consent to participate

Not applicable.

## Supplementary Information

**Table S1:** Genome annotation of phiE1336.

| Gene | Start | Stop | Strand | Annotation | Domain |
| --- | --- | --- | --- | --- | --- |
| gp1 | 1 | 2469 | + | Phage tail protein | - |
| gp2 | 2595 | 3632 | + | Phage tail protein | - |
| gp3 | 3654 | 3821 | + | HP | - |
| gp4 | 3825 | 4187 | + | HP | - |
| gp5 | 4376 | 5575 | + | LysinA | Amidase/PGRP |
| gp6 | 6065 | 5772 | - | HP | - |
| gp7 | 6081 | 6191 | + | HP | - |
| gp8 | 6188 | 7147 | + | LysinB_MF | PG binding and<br>AB_hydrolase |
| gp9 | 7147 | 7236 | + | HP | - |
| gp10 | 7242 | 7457 | + | Holin | - |
| gp11 | 7454 | 7900 | + | HP | - |
| gp12 | 8509 | 8093 | - | HP | - |
| gp13 | 8899 | 8513 | - | HP | - |

**Figure S1:**
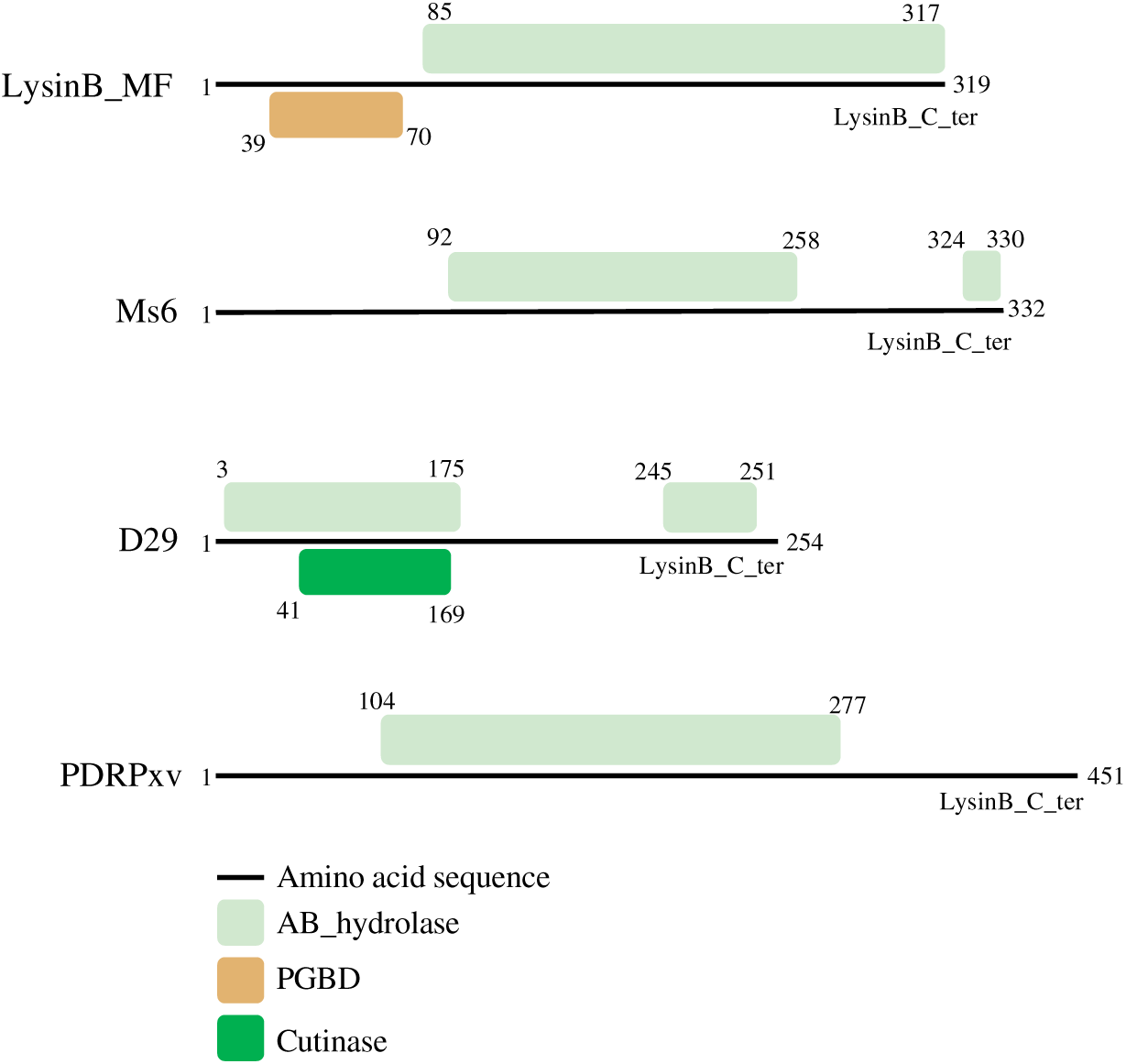
Comparison of domain organization of LysinBs. Domain prediction and architecture of well-studied LysinBs indicate the modular nature of LysinB_MF, containing a PGBD in the N-terminal end of the enzyme.

**Figure S2:**
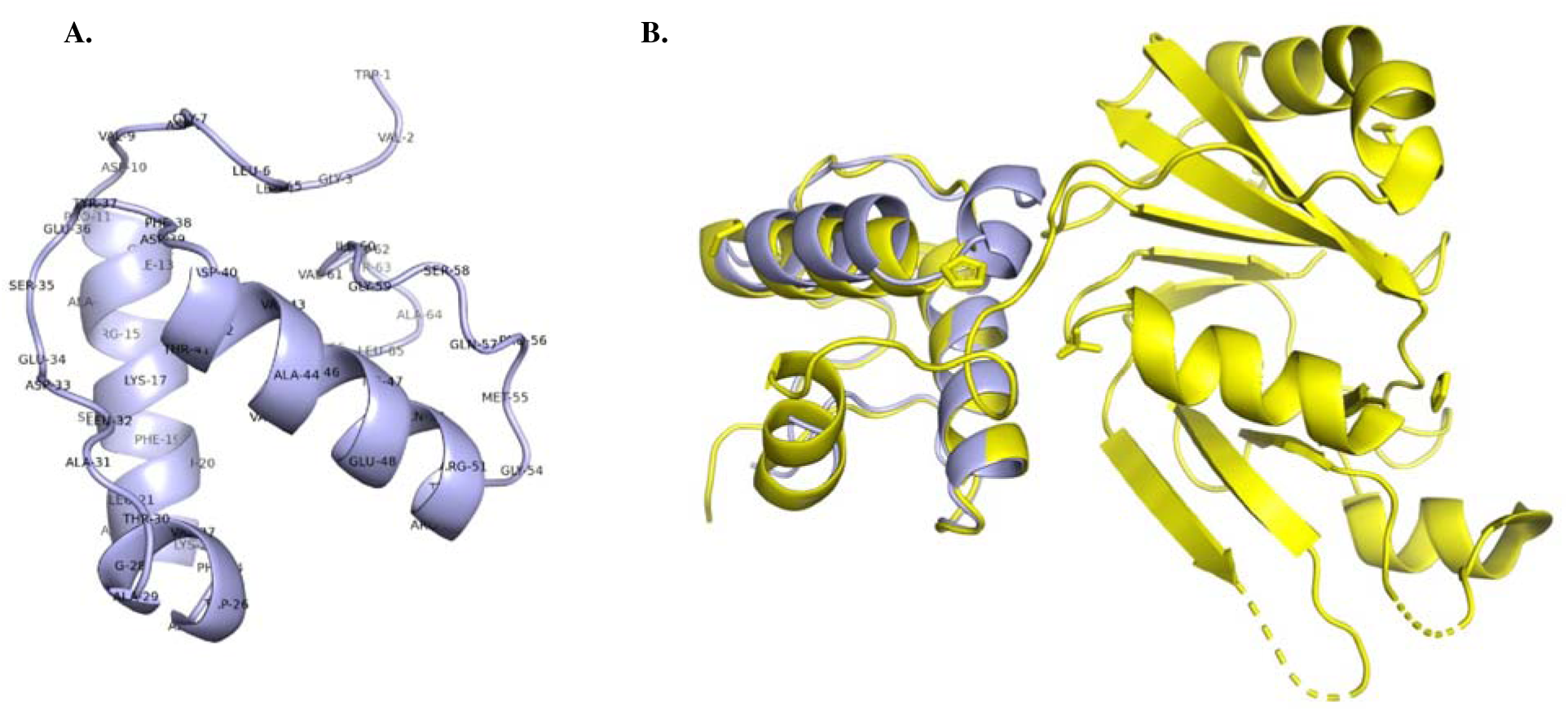
Structural prediction of PGBD from LysinB_MF and alignment with an amidase. **A)** To facilitate our understanding of the structural similarity of PGBD from LysinB_MF, we separately built its 3D model using the structure of the lysin as a template. The structure retains its original lz-helices, as observed in the native LysinB_MF. However, the single β-strand is missing. The structure shows very high confidence and was taken forward for further analysis. B) A zinc-dependent amidase (PDB accession 4XXT), a peptidoglycan-binding domain-containing protein from *Clostridium acetobutylicum* (ATCC 824), displays high structural similarity with the PGBD from LysinB_MF. The two structures can be superimposed and superpositioned with an overall RMSD as low as 0.46.

